# Active gelation breaks time-reversal-symmetry of mitotic chromosome mechanics

**DOI:** 10.1101/296566

**Authors:** Matthäus Mittasch, Anatol W. Fritsch, Michael Nestler, Juan M. Iglesias-Artola, Kaushikaram Subramanian, Heike Petzold, Mrityunjoy Kar, Axel Voigt, Moritz Kreysing

## Abstract

In cell division, mitosis is the phase in which duplicated sets of chromosomes are mechanically aligned to form the metaphase plate before being segregated in two daughter cells. Irreversibility is a hallmark of this process, despite the fundamental laws of Newtonian mechanics being time symmetric.

Here we show experimentally that mitotic chromosomes receive the arrow of time by time-reversal-symmetry breaking of the underlying mechanics in prometaphase. By optically inducing hydrodynamic flows within prophase nuclei, we find that duplicated chromatid pairs initially form a fluid suspension in the nucleoplasm: although showing little motion on their own, condensed chromosomes are free to move through the nucleus in a time-reversible manner. Actively probing chromosome mobility further in time, we find that this viscous suspension of chromatin transitions into a gel after nuclear breakdown. This gel state, in which chromosomes cannot be moved by flows, persists even when chromosomes start moving to form the metaphase plate. Complemented by minimal reconstitution experiments, our active intra-nuclear micro-rheology reveals time-reversal-symmetry breaking of chromosome mechanics to be caused by the transition from a purely fluid suspension into an active gel.

**Graphical abstract:** 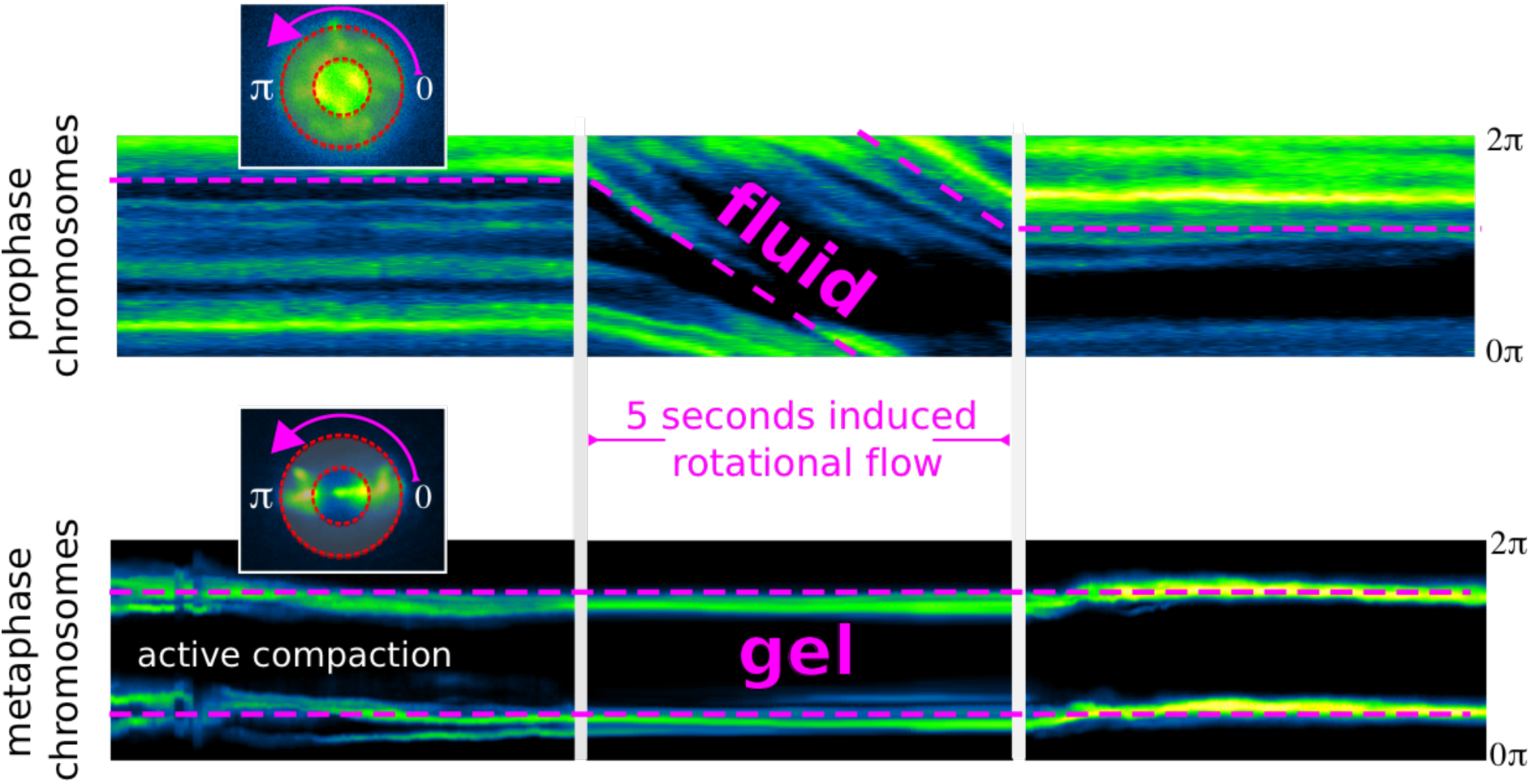

**One sentence summary:** Flows induced in living cell nuclei reveal the rheological changes that bring chromosomes under mechanical control during mitosis.

## Introduction

Biology is forward directed in time. Cells divide (Fig. 1a), but rarely fuse (Fig. 1b). A key process characteristic of this temporal directionality is mitosis (Fig. 1c), during which replicated sister chromatids are mechanically separated by the mitotic spindle apparatus and segregated into the two emerging daughter cells. The reverse, the fusion of two daughter cells into one cell, does usually not happen.

**Figure 1:**
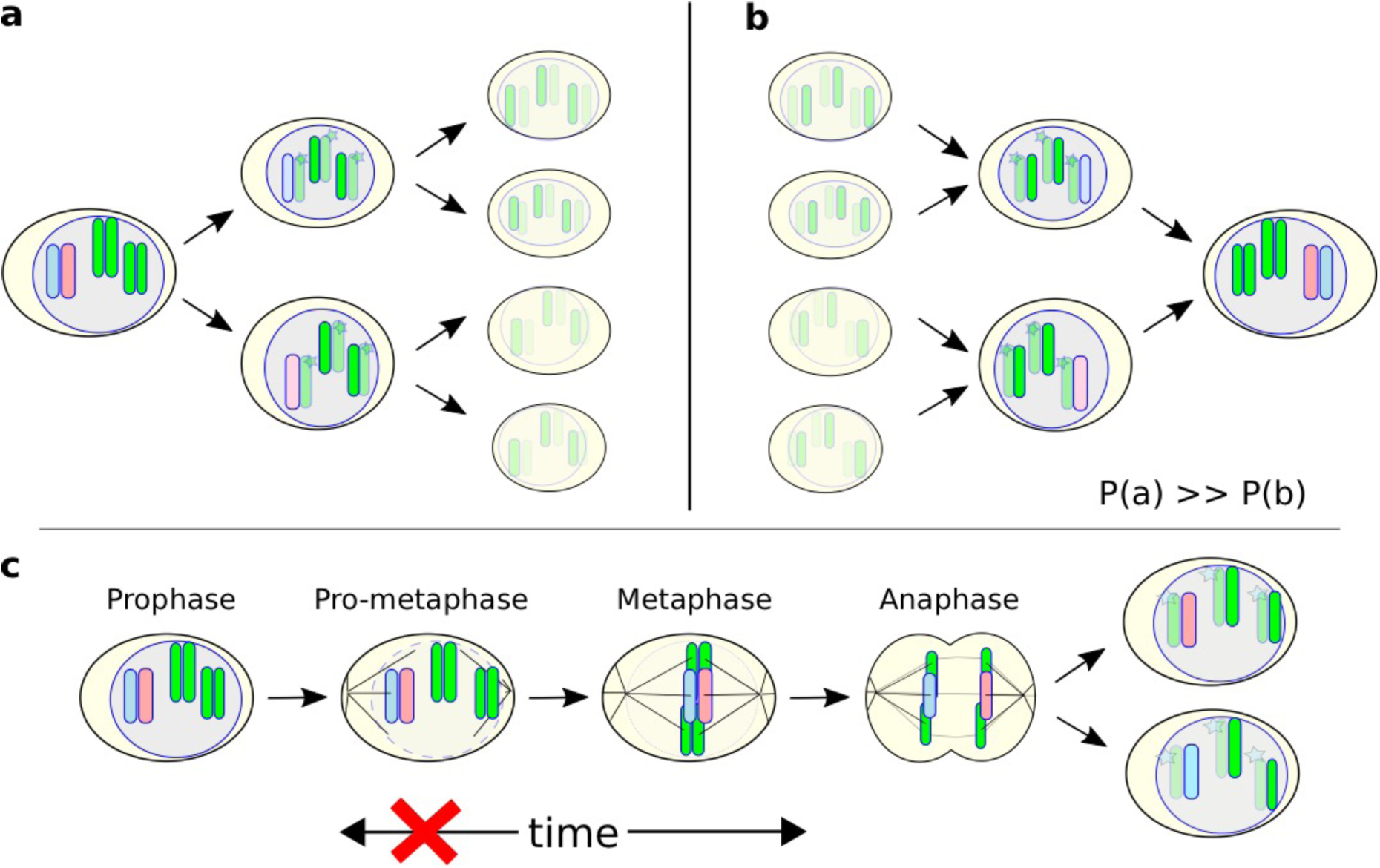
Mitotic chromosomes disobeys time symmetry. **a**, Cells commonly proliferate by division. **b**, In stark contrast, the time-reversed process, the sequential fusion of the same cells, is rarely observed. This raises the question of how biological cells break time-reversal-symmetry of Newtonian mechanics during mitosis, such that **c**, duplicated chromosomes may irreversibly be segregated from the mother cell into two daughter cells.

It seems both intuitive and sensible that cells divide in forward direction ^1,2^. And yet, the fundamental laws of mechanics, and their extension into the quantum world are time symmetric ^3,4^, meaning they do not have the arrow of time built in. Therefore, the questions emerge i) when and ii) how cells break time-reversal-symmetry of chromosome mechanics, such that chromosome segregation becomes a forward directed and irreversible process.

Herein, we built on recent methodological advancement that allows us to optically induce hydrodynamic flows within living cells and developing embryos ^5,6^, a technique we refer to as focused-light-induced cytoplasmic streaming (FLUCS). Physically, FLUCS makes use of thermoviscous flows ^7^, which have been also used to distinguish between fluid and gel like states of the cytoplasm in yeast cells ^5^.

## Results

Using FLUCS, we experimentally find and understand through simulations ^5,8^ that hydrodynamic flows can also be introduced on nuclear length scales (Fig. 2a-d, Supplementary Movie 1), and indeed in mitotic cell nuclei (Fig. 2e-f, Supplementary Movie 2-4). When inducing flows in prophase cell nuclei (*C. elegans* embryo 1-16 cell stage) we find that flows causing a full 180-degree rotation of chromosome position can be induced at strictly physiological conditions, meaning at temperature changes down to ±1 *Kelvin*.

**Figure 2:**
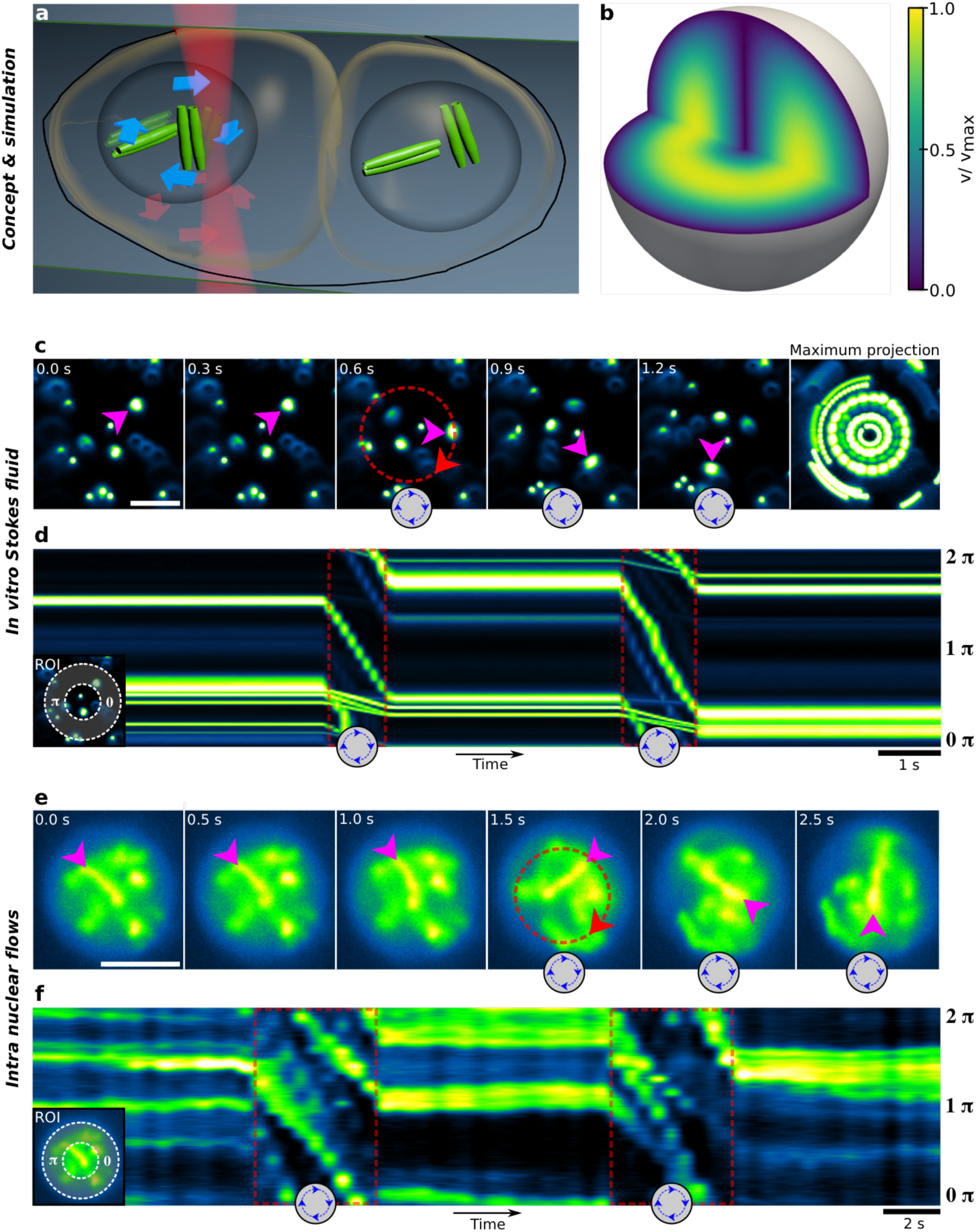
Prophase chromosomes are suspended in a Stokes fluid. **a,** Schematic of directed flows optically induced in the nucleoplasm of developing *C. elegans* embryos. **b,** Finite element modelling showing 3d profile of flows induced in these cell nuclei. **c,** Flows induced in highly viscous colloidal suspension (sucrose based), **d,** kymograph analysis in indicated region of interest (ROI), showing the instantaneous colloidal motion with the induced flow. (e) Chromosome rotational motion in response to induced flows in 2 cell *C. elegans* embryo (*dT* < ±2 *k*, fluorescence H2B-eGFP). **e,** Kymograph of induced chromosome translocation reveals near identical response to (c-d) meaning, i) little chromosome motion prior to flows, ii) instantaneous motion of chromosomes with induced flows, and iii) the immediate, creep-free arrest of chromosomal motion after flow stimulus.

We will show that this active induction of flows in nuclei of living cells enables us to infer mechanical properties of chromosome mechanics beyond passive micro-rheology ^9^. By kymograph analysis, we find that chromosomes suspended in prophase nuclei (Fig. 1f) behave near identical to colloids in a pure Stokes fluid, i.e. a highly viscous sucrose solution (Fig. 1d). Specifically, chromosome motion starts and stops immediately with the induction of flows. This instantaneous behavior demonstrates i) the absence of inertia, and ii) excludes inter-chromosomal elastic interactions that would cause creep relaxation after flow inductions. To directly compare flow-driven transport with diffusive motion of the chromosomes, we calculate a Péclet number of *pe* ≥ *Lv*/*D* = 40 (nuclear diameter of *L* = 10 µ*m*, a typical velocity of *v* = 1 µ*ms*^−1^ and an experimentally determined diffusion coefficient of *D* < 0.25 µ*m*^2^*s*^−1^, see supplement) specifying the strongly directed character of chromosome motion when subjected to FLUCS.

Our active measurements suggest that prophase chromosomes form a viscous suspension in a Stokes-fluid-like nucleoplasm. In order to better understand if chromosomes move collectively, or individually we use non-rotationally symmetric flow stimuli that translate into non-rotationally symmetric flow fields (Fig. 3a, Supplementary Notes), and analyzed chromosome motion in a co-rotating frame of reference (Fig. 3b). By this, we find that induced flows are able to interchange the relative positioning of chromosomes, even at later developmental stages (Fig. 3c-d). This ability to exchange chromosome neighbors will have strong implications to better understand structure-function relationship of nuclear architecture ^10-18^. In the course of this study it should be highlighted that individual prophase chromosomes, rather than large clusters, form a colloidal suspension inside a Stokes fluid.

**Figure 3:**
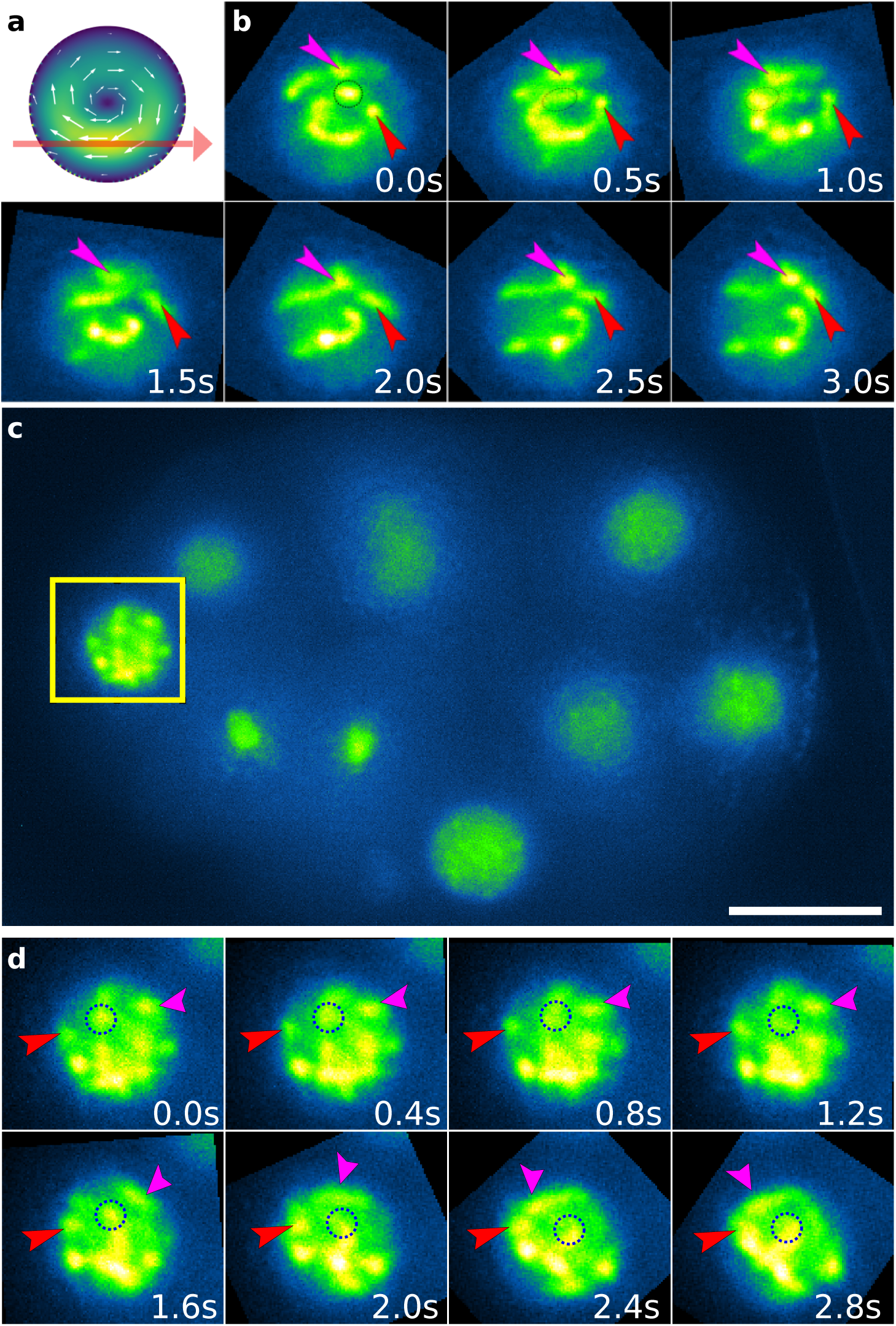
Prophase chromosomes are free to exchange neighbors. **a,** Simulation: the asymmetric laser scanning over the nucleus induces non-rotationally symmetric flow fields. **b,** Experiment: the flow-induced re-arrangement of prophase chromosomes is shown in co-rotating frame of reference for two cell *C. elegans* embryo. Arrows highlight relative positioning of two chromosomes. **c,** Later embryo referencing nucleus in **(d):** co-rotating frame of reference. Two chromosomes (arrows) are initial separated by a third one (circle). When flows are introduced the previously separated chromosomes become neighbors. Scale bar 10 µm.

We next estimated the magnitude of flow induced drag forces in the nucleus. These can well calculated using the Stokes-Einstein relation ^19^. Using *K*_b_T as thermal energy at 20°C, an experimentally determined diffusion constant of chromosomes *D* = 0.25 µ*m*^2^*s*^−1^ (which is in good agreement with colloids of 2 µ*m* diameter in water *D*_*2µm*_ = *K*_b_*T*/(6πη*r*) = 0.239 µ*m*^2^*s*^−1^ and *v* = 1µ*ms*^−1^, we calculate a drag force of *F* = *K*_b_T/*Dv* = 16 *fNewton*, which is 5 orders of magnitude lower than the force required to break a single covalent bond (1.4 − 2.0 *nN*, ^20^). This force comparison implies i) that chromosomes dragged with the flow must be freely suspended to follow it, ii) stresses the non-invasive character of the perturbation, and suggests iii) that mechanical changes of this chromosome suspension should be detectable as discrete events with single molecule sensitivity.

Following the behavior of a Stokes fluid, chromosome mechanics in the prophase nucleus should obey time-reversal-symmetry. While time-reversal-symmetry is difficult to observe in passive systems that receive their dynamics from diffusion only, active mechanical perturbations of these systems can be well suited to show time-reversible or periodic transitions from one mesoscopic state into another. Famous examples of this include the reversible shearing of localized dye or colloidal particles in a Couette cylinder. ^21-25^

Here we use FLUCS in order to directly test if the suspension of chromosomes in the prophase nucleus obey time-reversal-symmetry. For this we applied time-symmetric flow stimuli in nuclei of two cell *C. elegans* embryos. These flow stimuli start with a clockwise rotational motion of the nucleoplasm, followed by the precisely inverted, counterclockwise stimulus of the same duration and velocity (Fig. 4a, Supplementary Movie 5). As a result, we find that chromosomes are moved within 2 seconds from state *X*(−2*s*) to state *X*(0), which shows little resemblance with the original state *X*(−2*s*). Upon reversal of velocities at t = 0s, chromosomes move to state *X*(2*s*), which closely resembles the original state *X*(−2*s*). Hence, the reversal of velocities *v* → −*v* produces the same result as the reversal of time *t* → −*t*. This fingerprint of time-symmetric mechanics can be seen even more clearly when continuing this oscillation, and plotting the systems response as a kymograph (Fig. 4b, Supplementary Movie 6). This kymograph directly reveals mirror axes of time to be located at the turning points of the periodic motion (Fig. 4b). Minor deviations from perfectly symmetric trajectories are due to residual diffusive motion, which can even be observed for colloids immersed in a highly viscous sugar solution (Fig. 4c). We conclude that despite the thermodynamic irreversibility of cellular biochemistry, prophase chromosomes constitute a mechanical system that obeys time-reversal-symmetry.

**Figure 4:**
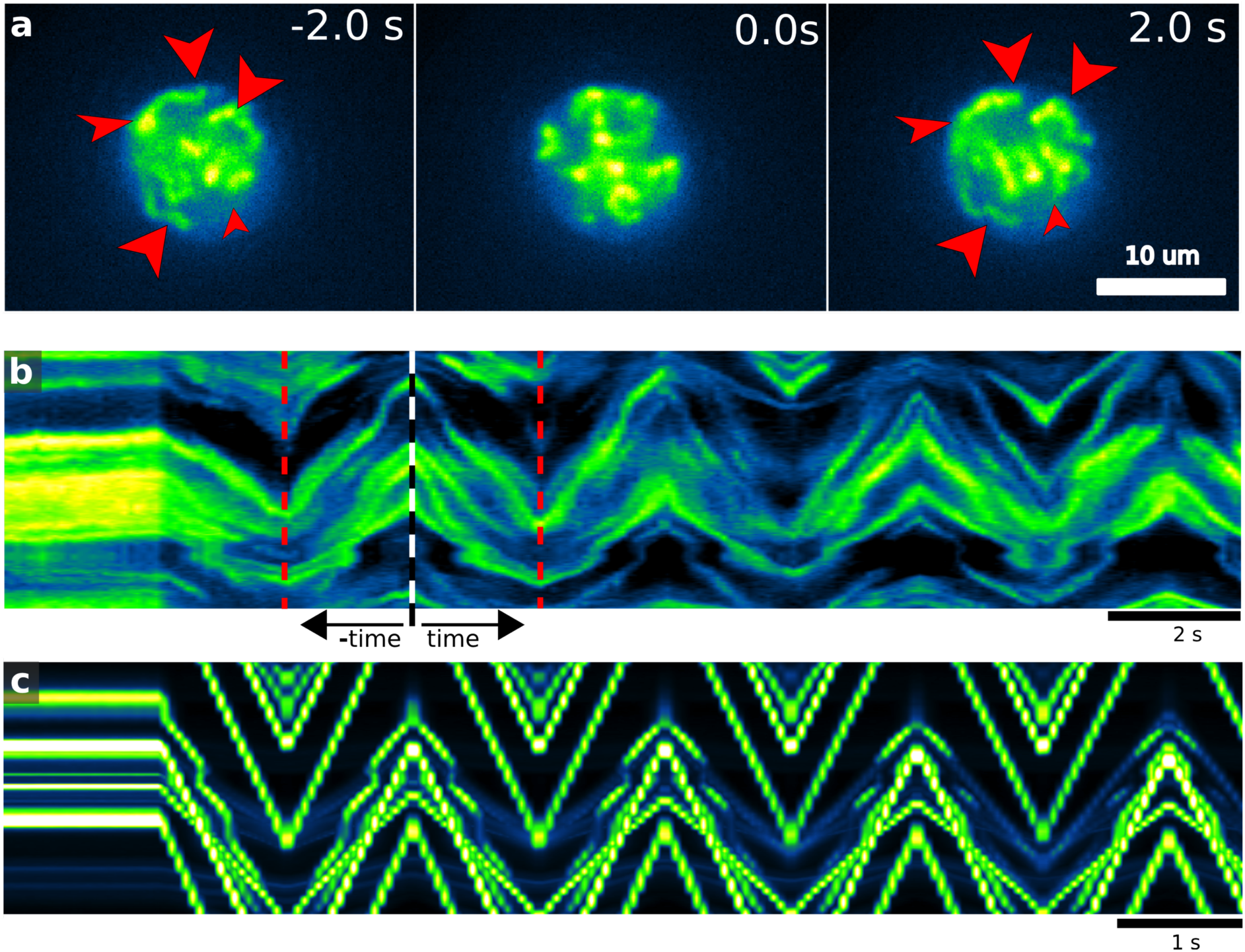
Prophase chromosome mechanics obey time-reversal-symmetry. **a,** Time-symmetric flow stimuli yield time-symmetric chromosome trajectories in a two cell *C. embryo*. **b,** Symmetrically alternating flow stimuli: reversal of flows after a perturbation yields the same results as the reversal of time. **c,** Reconstituted system of colloids suspended in Stokes fluid shares time-symmetric behavior of prophase chromatin.

Mitosis, however, is an irreversible process, which becomes apparent when initially freely suspended chromosomes start congressing to form the metaphase plate (Fig. 5a, b, Supplementary Movie 7). In physical terms, the active nature^26^ of this process can be observed as an entropy decrease of this colloidal system (Fig. 5c). This old observation of directed chromosome motion^27-30^ emphasizes that the time-symmetric mechanics of prophase chromosomes motion must fundamentally change, such that mitosis may become a process forward-directed in time.

**Figure 5:**
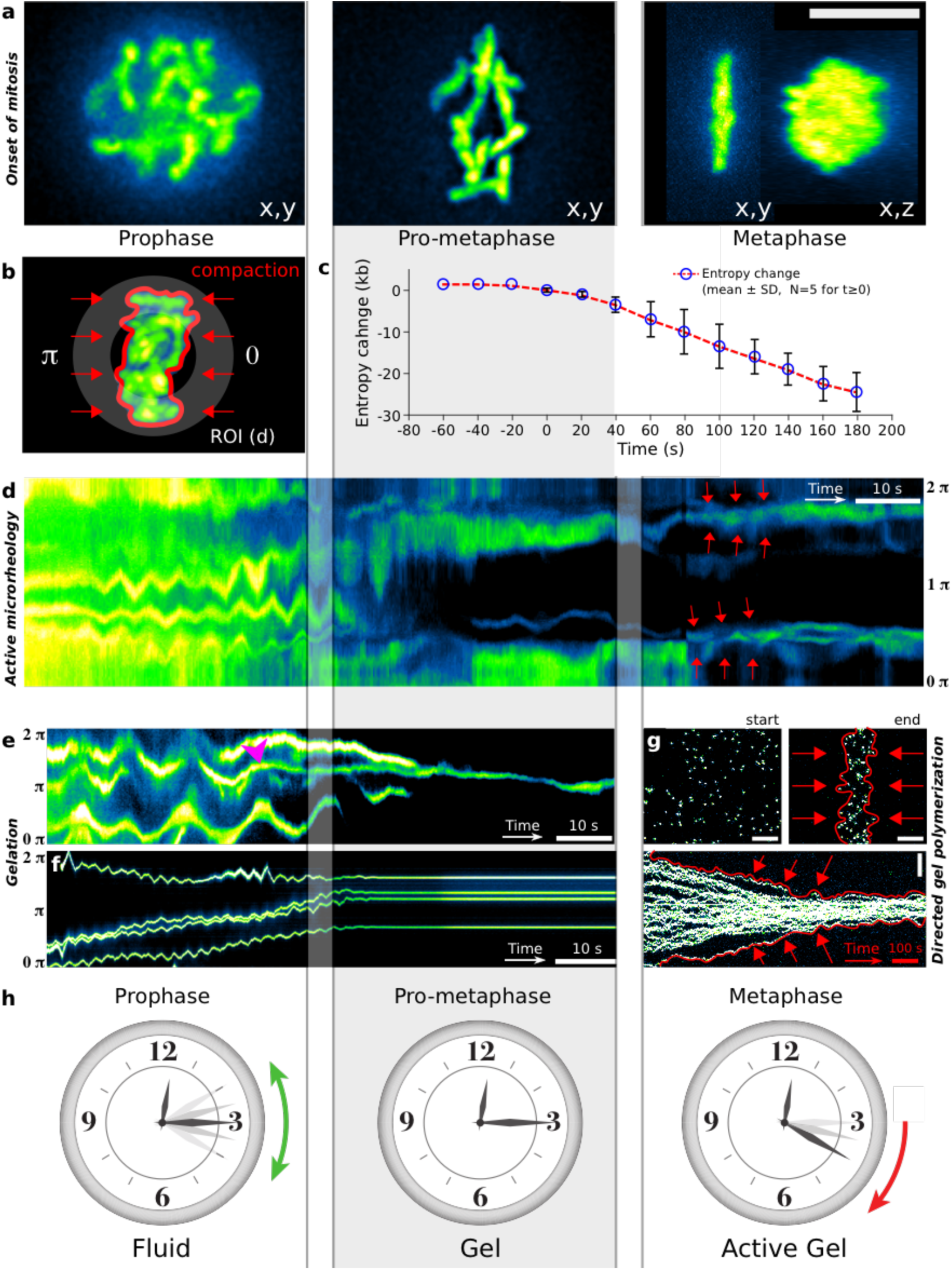
Active gelation breaks time reversal-symmetry of mitotic chromosomes. **a,** Mitotic phases characterized by active flow rheology. **b**, Volume estimation and region of interest for c and d respectively. **c**, Compaction-induced entropy reduction in colloidal system of chromosomes as a fingerprint of active processes at work. **d,** Kymograph of chromosome oscillations showing two transitions of the mechanical state: i) a fluid-to-gel transition after nuclear envelope breakdown, which renders chromosome fully resistible to induced flows. ii) The transition into an *active gel*, that starts to move and compacts (red arrows), although it cannot be moved by flows anymore. **e**, Detailed analysis of the gel transition shows that individual chromosomes stop moving suddenly (pink arrow), while other ones keep on oscillating for multiple periods nearby. **Sufficiency: f,** Gelation causes mobility arrest of polyacrylamide suspended colloids. **g,** Directed gel polymerization (red arrows) suffices to cause irreversibly directed motion of colloids, and their compaction in metaphase plate type geometry. **h**, Physical model of mitotic chromosome mechanics: prophase chromosomes constitute a Stokes fluid-type suspension for which positive and negative time are equivalent. In pro-metaphase this fluid phase transitions into a gel, which is dominated by elastic forces. Time-reversal-symmetry is broken when this gel becomes active and directionally moves chromosomes, although they cannot be moved by induced flows anymore.

In order to monitor the mechanics of chromosomes beyond prophase, we applied small amplitude oscillatory flow stimuli (*f* = 0.25 *Hz*) as the cell cycle progressed (Fig. 5d, Supplementary Movie 8). Due to the active nature of induced flows, our measurement allows us to infer changes in mechanical properties also where passive micro-rheology is limited due absence of spontaneously occurring motions. Our flow oscillation data strikingly shows that upon transition into pro-metaphase, shortly after the nuclear envelope break-down, the previously observed periodic motion of chromosomes comes to a halt (Fig. 5d center). Now chromosomes persist in their positions despite the flow driving stimuli being still present. When investigating this transition in more detail, we find that individual chromosomes stop moving suddenly (Fig. 5e, pink arrow), while others near by may be oscillating for several subsequent periods. These discrete motion arrests evidence sudden local increase in elastic restoring forces, rather than the cease of oscillation driven hydrodynamic flows. The fact that drag forces on individual chromosome are 4-5 orders of magnitude smaller than the rupture force of a single covalent bond suggests that the discrete motion arrest of individual chromosomes constitutes their binding to individual microtubules. This low magnitude of flow induced shear forces makes FLUCS a particularly sensitive read out of gelation processes, and less invasive than previous approaches of chromosome repositioning using glass needles^31^.

The mechanical properties of the viscous suspension of prophase chromosomes thus seems to undergo a step-wise discrete transition from a Stokes Fluid into a gel state, which resists flow-induced drag forces. In support of this gelation model, we observe a sudden resistance to flows also when inducing the gelation of a monomeric Stokes Fluid by photo-activation chemistry ^32^ (Fig. 5f, see Supplement for details). As we further show, it is the transition into this gel state that initiates the breaking of time-reversal-symmetry to define the arrow of time of chromosome mechanics during mitosis.

When progressing in the cell cycle, through pro-metaphase and towards metaphase, chromosomes are known to congress to the nuclear center ^33,34^ to form the metaphase plate (Fig. 5a, r.h.s). By active flows rheology measurements, we find that despite this spindle-mediated increase in mobility ^35^, congressing chromosomes keep on resisting induced flows (Fig. 5d, r.h.s) which indicates their persistent association to a gel-like matrix. Contrasted by their initial fluid state, in which prophase chromosomes showed little spontaneous motion despite being free to move, this directed motion towards the metaphase plate seems incompatible with Einstein’s formulation of the Fluctuation-Dissipation theorem. The theorem centrally states that under equilibrium conditions the spontaneous motion of colloidal particles requires their passive mobility ^36^.

And indeed, when investigating mobility changes more systematically, we find that the same 5 second lasting flow stimuli that induces more than a full rotation of chromosomes during prophase (*v*_max_ > 4µ*ms*^−1^), moves metaphase chromosomes less than 500nm (Supplementary Figure 1 & Movie 9). These perturbations indicate an elastic restoring potential during metaphase in excess of *k* = (*vK*_b_T/*D*)/*d* > 120*fNµm*^−1^ per chromosome pair, which renders their thermal fluctuations of more than 260 *nm* implausible.

To explain this violation of the Fluctuation-Dissipation theorem, we suggest that long-range chromosome congression to the metaphase plate, which happens while chromosomes cannot be moved by flows, must be interpreted as the activity of a gel^37^. Growing inwards from two poles towards the mid-plane, this active gel state mediates high resistance to flow-induced frictional forces, and maintains tight mechanical control over chromosome positioning. Apart from violating the Fluctuation-Dissipation theorem, active gels can also break time-reversal-symmetry of Newtonian mechanics ^37^: We furthermore explicitly show by minimal *in vitro* experiments that time-reversal-symmetry breaking of colloidal suspensions can be achieved by directed polymerization, reminiscent of spindle growth within the cell. When polymerized from two opposing poles, a growing polyacrylamide gel induces irreversible directed colloidal motion and ultimately compacts micrometer-sized beads in a mid-plate type configuration (Fig. 5g, Supplementary Movie 10). Hence, the directed polymerization of a gel suffices to lower entropy in colloidal system by compaction.

## Discussion

While cells are well-known to operate in a thermodynamically irreversible manner, subcellular systems can still obey time-reversal-symmetry. We demonstrated this for prophase chromosomes, which we discovered to represent a fluid suspension.

Due to the active nature of our micro-rheology measurements, we could further show that chromosomes transition from a purely fluid suspension, to become part of a rigid gel (Fig. 5, S1) during early mitosis. Remarkably, this gel-state still persists when chromosomes are actively moved to form the metaphase plate. In light of our new experiments, assumptions implicit to models of chromosome centering by *polar winds* or chromosome pushing forces should be reconsidered ^38-40^. Our experiments clearly show that in *C. elegans* embryos, forces exerted by the spindle do not primarily serve to compensate drag forces of chromosomes congressing in a viscous medium. If this was the case, viscous (dissipative) forces would cause significant fluctuations when flows are optically induced. These flow-induced oscillations, however, reduce to barely detectable amplitudes during the active process of metaphase plate formation (Fig. S1), stressing that while chromosomes are dynamically aligned, they are part of a rigid gel.

## Conclusion

Our active flow rheology measurements show that mitotic chromatin mechanics receives a forward-directed arrow of time in a two-step process: first, discrete binding events cause a transition from a purely viscous suspension of chromosomes, to a predominantly elastic gel. Second, it is the activity of this persistent gel-state that beaks time-reversal-symmetry, thereby tightly guiding chromosome congression in perturbation-resistant and mechanically irreversible manner. Beyond the mesoscale characterization of chromosomes as physical phase, the here presented ability to alter chromosome positioning will leverage our understanding of the structure-function-relationship of nuclear architecture also beyond mitosis^11-17^.

## Author Contributions

MM, AF, MK: carried out FLUCS experiments and analyzed data, MN & AV performed computer simulations, MM & AF built microscope, MKa, JI & MK carried out gel reconstitution experiments, AF, JI and KS acquired and analyzed video microscopy data, HP cultured worms, MK conceived project, and wrote the paper. All authors critically discussed the data and the final manuscript.

## Acknowledgements

The authors acknowledge support from the Max Planck Society, a DFG-financed DIPP fellowship for M.Mi., a Boehringer Ingelheim Fonds PhD fellowship for JI, infrastructural support and worms by the Hyman lab, and technical support from MPI-CBG light-microscopy and scientific computing facilities. The authors thank J. Brugués sharing computational code for diffusion analysis, P. Tomancak, F. Jülicher, A. Hyman, V. Zaburdaev, É. Roldán, C. Hoege, Stefan Grill, M. Weigert, S. Bundschuh (all Dresden) and Ulrich Schwarz (RKU Heidelberg) for helpful discussions and comments.

## Supplementary Information

**Supplementary Figure 1:**
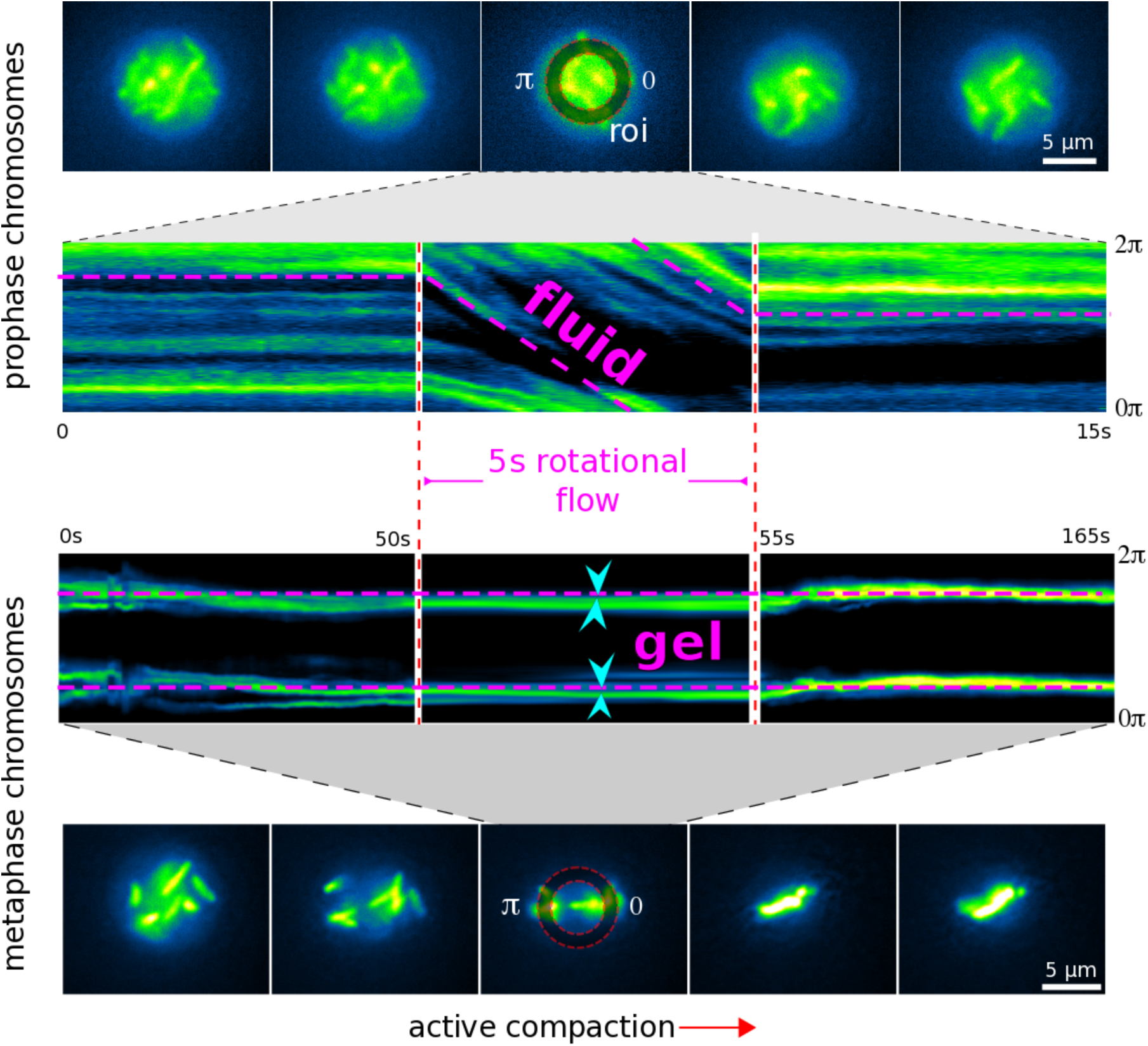
Differential mobility of prophase *versus* metaphase chromosomes. The same 5 second flow-inducing stimulus that causes a full rotation of chromosomes during prophase, displaces chromosomes typically less than 500nm during metaphase, indicating the dominance of elastic restoring forces present in a gel.

## Description of supplemental videos

**SMovie 1**: Thermoviscous flows in a viscous colloidal suspension.

Rotational flows are introduced on nuclear length scales. The Stokes fluid is represented by a high concentration sucrose solution in which fluorescent beads are immersed. Length of movie is 10 seconds.

**SMovie 2**: Flows induced in embryonic *C. elegans* nuclei move condensed chromosomes. Rotational flows are induced in a prophase nucleus of a 2 cell C. elegans embryo. Flows share characteristics of Stokes flows as they respond instantaneously to the flow stimulus and show no inertia (low Reynolds number regime), or elastic restoring forces after rotation.

**SMovie 3**: Flows induced in nuclei of 4 cell *C. elegans* embryos.

**SMovie 4**: Flows induced in nuclei of 12 cell *C. elegans* embryos. Clockwise flows are introduced in l.h.s. cell starting from 5.5 sec, followed by counter-clockwise flows.

**SMovie 5**: Prophase chromosomes show time symmetric motion in response to a time symmetric flows stimulus. A clockwise flow stimulus is reversed after 2 seconds of application. Another 2 seconds later, the chromosomes returned to their initial state in time symmetric manner. Full length of video 4 seconds.

**SMovie 6**: Prophase chromosomes show time symmetric motion in response to a periodic flow stimulus.

**SMovie 7**: Prophase to metaphase transition of chromosomes in *C. elegans* embryos, shows increased chromosome mobility and compaction during metaphase plate formation. Confocal microscopy data, 2 orthogonal views.

**SMovie 8**: Flow driven microrheology during early mitosis. Oscillatory flows move chromosomes during prophase. This oscillation come to a hold when chromosomes congress to form the metaphase plate.

**SMovie 9**: Differential mobility of prophase vs metaphase chromosomes.

The same flow inducing stimulus that induces a full rotation of chromosomes during prophase, has no visible effect on chromosomes during metaphase, indicating the dominance of elastic restoring forces present in a gel.

**SMovie 10**: Directed polymerization aligns colloids. Polymerization of a polyacrylamide gel from the vertical margins aligns suspended colloids in a metaphase plate type geometry. Duration of video is 12 minutes.

## References

1. Eigen, M. From Strange Simplicity to Complex Familiarity. (OUP Oxford, 2013). doi:10.1093/acprof:oso/9780198570219.001.0001

2. Schwille, P. Bacterial Cell Division: A Swirling Ring to Rule Them All? Current Biology 24, R157–R159 (2014).

3. Newton, I. The Principia. (University of California Press, 1999).

4. Schrödinger, E. An Undulatory Theory of the Mechanics of Atoms and Molecules. Phys. Rev. 28, 1049–1070 (1926).

5. Mittasch, M. et al. Non-invasive perturbations of intracellular flow reveal physical principles of cell organization. Nature Cell Biology 2018 20:3 20, 344–351 (2018).

6. Mittasch, M. et al. How to apply FLUCS in single cells and living embryos. (2018). doi:10.1038/protex.2017.157

7. Weinert, F. M. & Braun, D. Optically driven fluid flow along arbitrary microscale patterns using thermoviscous expansion. Journal of Applied Physics 104, 104701 (2008).

8. Nitschke, I., Voigt, A. & Wensch, J. A finite element approach to incompressible two-phase flow on manifolds. Journal of Fluid Mechanics 708, 418–438 (2012).

9. Ahmed, W. W., Fodor, É. & Betz, T. Active cell mechanics: Measurement and theory. Biochim. Biophys. Acta 1853, 3083–3094 (2015).

10. Arai, R. et al. Reduction in chromosome mobility accompanies nuclear organization during early embryogenesis in Caenorhabditis elegans. Sci Rep 7, 3631 (2017).

11. Cremer, T. & Cremer, M. Chromosome Territories. Cold Spring Harb Perspect Biol 2, a003889–a003889 (2010).

12. Gerlich, D. et al. Global Chromosome Positions Are Transmitted through Mitosis in Mammalian Cells. Cell 112, 751–764 (2003).

13. Naumova, N. et al. Organization of the Mitotic Chromosome. Science 342, 948–953 (2013).

14. Solovei, I. et al. Nuclear architecture of rod photoreceptor cells adapts to vision in mammalian evolution. Cell 137, 356–368 (2009).

15. Solovei, I. et al. LBR and lamin A/C sequentially tether peripheral heterochromatin and inversely regulate differentiation. Cell 152, 584–598 (2013).

16. Lanctôt, C., Cheutin, T., Cremer, M., Cavalli, G. & Cremer, T. Dynamic genome architecture in the nuclear space: regulation of gene expression in three dimensions. Nature Reviews Genetics 2007 8:28, 104–115 (2007).

17. Falk, M. et al. Heterochromatin drives organization of conventional and inverted nuclei. bioRxiv 244038 (2018). doi:10.1101/244038

18. Bolzer, A. et al. Three-Dimensional Maps of All Chromosomes in Human Male Fibroblast Nuclei and Prometaphase Rosettes. PLOS Biology 3, e157 (2005).

19. Einstein, A. Über die von der molekularkinetischen Theorie der Wärme geforderte Bewegung von in ruhenden Flüssigkeiten suspendierten Teilchen. ANNALEN DER PHYSIK 322, 549–560 (1905).

20. Grandbois, M., Beyer, M., Rief, M., Clausen-Schaumann, H. & Gaub, H. E. How Strong Is a Covalent Bond? Science 283, 1727–1730 (1999).

21. Heller, J. P. An Unmixing Demonstration. American Journal of Physics 28, 348–353 (1960).

22. Eirich, F. R. Rheology: Theory and Applications. 4, (Academic Press, 1969).

23. Morrison, I. D. & Ross, S. Colloidal Dispersions: Suspensions, Emulsions, and Foams. (John Wiley & Sons, 2002).

24. Leal, L. G. Laminar Flow and Convective Transport Processes. Butterworth-Heinemann (1992).

25. Fonda, E. & Sreenivasan, K. R. Unmixing demonstration with a twist: A photochromic Taylor-Couette device. American Journal of Physics 85, 796–800 (2017).

26. Needleman, D. & Dogic, Z. Active matter at the interface between materials science and cell biology. Nature Reviews Materials 2017 2:9 2, 17048 (2017).

27. Berns, M. W. et al. Use of a laser-induced optical force trap to study chromosome movement on the mitotic spindle. PNAS 86, 4539–4543 (1989).

28. Brouhard, G. J. & Hunt, A. J. Microtubule movements on the arms of mitotic chromosomes: Polar ejection forces quantified in vitro. PNAS 102, 13903–13908 (2005).

29. Levesque, A. A. & Compton, D. A. The chromokinesin Kid is necessary for chromosome arm orientation and oscillation, but not congression, on mitotic spindles. The Journal of Cell Biology 154, 1135–1146 (2001).

30. Roldán, é., Neri, I., Dörpinghaus, M., Meyr, H. & Jülicher, F. Decision Making in the Arrow of Time. Phys. Rev. Lett. 115, 250602 (2015).

31. Nicklas, R. B. & Koch, C. A. CHROMOSOME MICROMANIPULATION: III. Spindle Fiber Tension and the Reorientation of Mal-Oriented Chromosomes. The Journal of Cell Biology 43, 40–50 (1969).

32. Dasgupta, B. R. & Weitz, D. A. Microrheology of cross-linked polyacrylamide networks. Phys. Rev. E 71, 021504 (2005).

33. Antonin, W. & Neumann, H. Chromosome condensation and decondensation during mitosis. Current Opinion in Cell Biology 40, 15–22 (2016).

34. Kapoor, T. M. et al. Chromosomes Can Congress to the Metaphase Plate Before Biorientation. Science 311, 388–391 (2006).

35. Begg, D. A. & Ellis, G. W. Micromanipulation studies of chromosome movement. I. Chromosome-spindle attachment and the mechanical properties of chromosomal spindle fibers. The Journal of Cell Biology 82, 528–541 (1979).

36. Mizuno, D., Tardin, C., Schmidt, C. F. & MacKintosh, F. C. Nonequilibrium Mechanics of Active Cytoskeletal Networks. Science 315, 370–373 (2007).

37. Prost, J., Jülicher, F. & Joanny, J.-F. Active gel physics. Nature Physics 2015 11:211, 111–117 (2015).

38. McIntosh, J. & Hays, T. A Brief History of Research on Mitotic Mechanisms. Biology 2016, Vol. 5, Page 55 5, 55 (2016).

39. Pease, D. C. Hydrostatic pressure effects upon the spindle figure and chromosome movement. I. Experiments on the first mitotic division of urechis eggs. Journal of Morphology 69, 405–441 (1941).

40. Pease, D. C. Hydrostatic Pressure Effects Upon The Spindle Figure And Chromosome Movement. Ii. Experiments On The Meiotic Divisions Of Tradescantia Pollen Mother Cells. The Biological Bulletin 91, 145–169 (1946).

